# Modular Phoneme Processing in Human Superior Temporal Gyrus

**DOI:** 10.1101/2024.01.17.576120

**Authors:** Daniel R. Cleary, Youngbin Tchoe, Andrew Bourhis, Charles W. Dickey, Brittany Stedelin, Mehran Ganji, Sang Hoen Lee, Jihwan Lee, Dominic A. Siler, Erik C. Brown, Burke Q. Rosen, Erik Kaestner, Jimmy C. Yang, Daniel J. Soper, Seunggu Jude Han, Angelique C. Paulk, Sydney S. Cash, Ahmed M. T. Raslan, Shadi A. Dayeh, Eric Halgren

## Abstract

Modular organization is fundamental to cortical processing, but its presence is human association cortex is unknown. We characterized phoneme processing with 128-1024 channel micro-arrays at 50-200μm pitch on superior temporal gyrus of 7 patients. High gamma responses were highly correlated within ∼1.7mm diameter modules, sharply delineated from adjacent modules with distinct time-courses and phoneme-selectivity. We suggest that receptive language cortex may be organized in discrete processing modules.

## Main Text

Cortical function is often understood as composed of discrete processing modules whose outputs converge on other modules. Commonly, cortical columns are posited to perform this role, bridging local circuits and cortical parcels. In primary sensory cortices of certain species, spatially demarcated groups of neurons form discrete columns that respond to specific stimuli, distinct from neighboring columns^1-3^. Using fMRI, orientation-selective and ocular dominance columns have been identified in V1^4^, color and binocular disparity stripes in V2 and V3^5^, axis of motion columns in MT^6^, and frequency columns in A1^7^. Column diameter varies. They are ∼500um in cat somatosensory^1^ or macaque ocular dominance, but human ocular dominance are ∼1.7x wider^8^. However, due to their spatial resolution, these fMRI studies have difficulty demonstrating a mosaic organization with functional properties relatively constant within the column and changing sharply at their boundaries. Furthermore, with the partial exceptions of MT and V2/3, these findings have been restricted to primary sensory cortex. Although columnar organization is clear anatomically^9,10^ it has not been associated with functional columns^11^. Thus, the status of columns, or other modularity at that scale, in human association cortex, remains unknown.

Direct intracranial recordings from the dominant posterior superior temporal gyrus (STG) have characterized the response correlates of neuronal populations to speech^12^. Generally, population firing is estimated from high gamma amplitude (HG; ∼70-150Hz), recorded from ∼3mm diameter contacts at 10mm pitch. Using 4mm pitch, Mesgarani et al^13^ found HG selectivity for groups of phonemes that shared articulatory features, and Leonard et al^14^ found that STG neurons in a track perpendicular to the surface often encode similar features, and their firing is reflected in the overlying surface. However, lateral spatial resolution in these studies has not been sufficient to resolve cortical columns.

Here, we measured the shape, size, and spacing of cortical modules selectively engaged by words using microgrids on the pial surface of STG. For the first set of experiments (4 participants), we used a 128-channel grid with 2×64 contacts at 50μm pitch (Fig. 1A). The second set of experiments (3 participants, one with 2 placements) used a 1024-channel 16×64 grid at 200μm pitch (Fig. 1B). All participants were undergoing acute awake craniotomies for mapping eloquent cortex prior to removal of a tumor^15^. Arrays were placed under visual control on a relatively avascular cortical surface which previous clinical testing identified as eloquent (Fig. 1C-D). Patients listened to Consonant-Vowel syllables and detected those that formed valid words. Noise-vocoding of the same stimuli produced controls with the same amplitude envelope in different frequency bands as the spoken stimuli (Fig. 1E).

**Fig 1.**
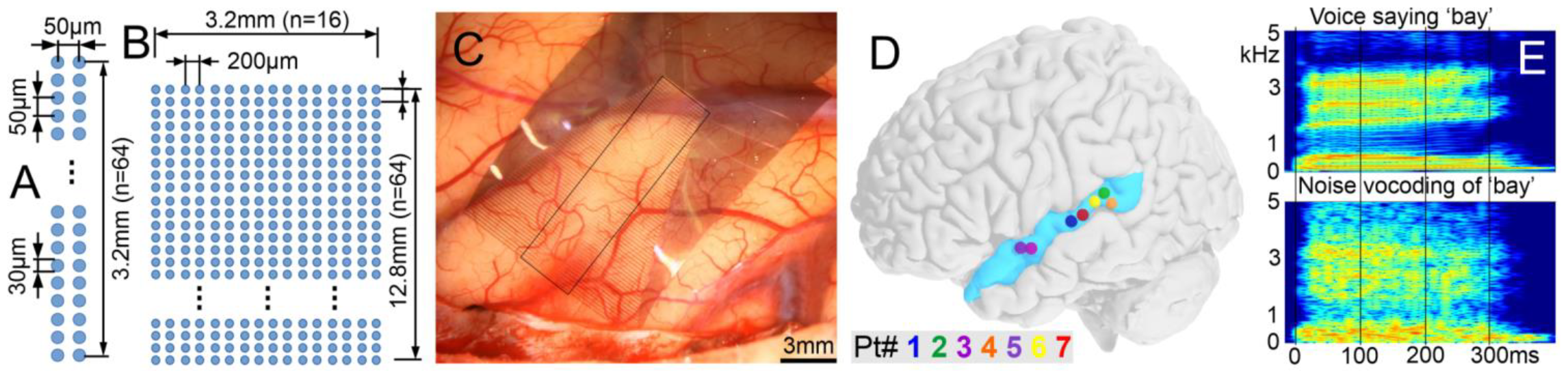
Electrodes and stimuli used in the preliminary experiments. Layouts of **A**. 128 ch array (2×64, 50µm pitch, 30μm diameter PEDOT contacts), and **B**. 1024 ch array (16×64, 200µm pitch, 30μm diameter Pt nanorod contacts). **C**. Intraoperative photo of 1024 ch array adhering to the pial surface. **D**. Electrode locations on standard brain. **E**. Spectrograms of an example word and its noise-vocoded control.

Six of the 7 patients showed significant modulation of local field potentials to words versus noise, in both lower frequencies (as the average evoked potential, Fig. 2A) and as the average instantaneous high gamma amplitude (HG; Fig. 2C). In the linear 2×64 arrays, both the evoked potential and HG responses were very similar for ∼25 channels (∼1.3mm) and then abruptly transitioned to channels with distinct response properties which again were similar to each other (Figs. 2BD). Within these cortical ‘modules,’ gamma-band correlations plateaued at a very high level, but at the boundary between modules, correlations dropped rapidly (Fig. 2F). The consistency of responses within modules and the steep transition between modules was also apparent in other measures, including the peak value of the HG to Words or Noise at each location (Fig. 2G). These empirical curves found a transition zone between modules of 400μm, which corresponds to that which would be generated by a disc of dipoles 100μm below the cortical surface (Fig. 2H).

**Fig 2.**
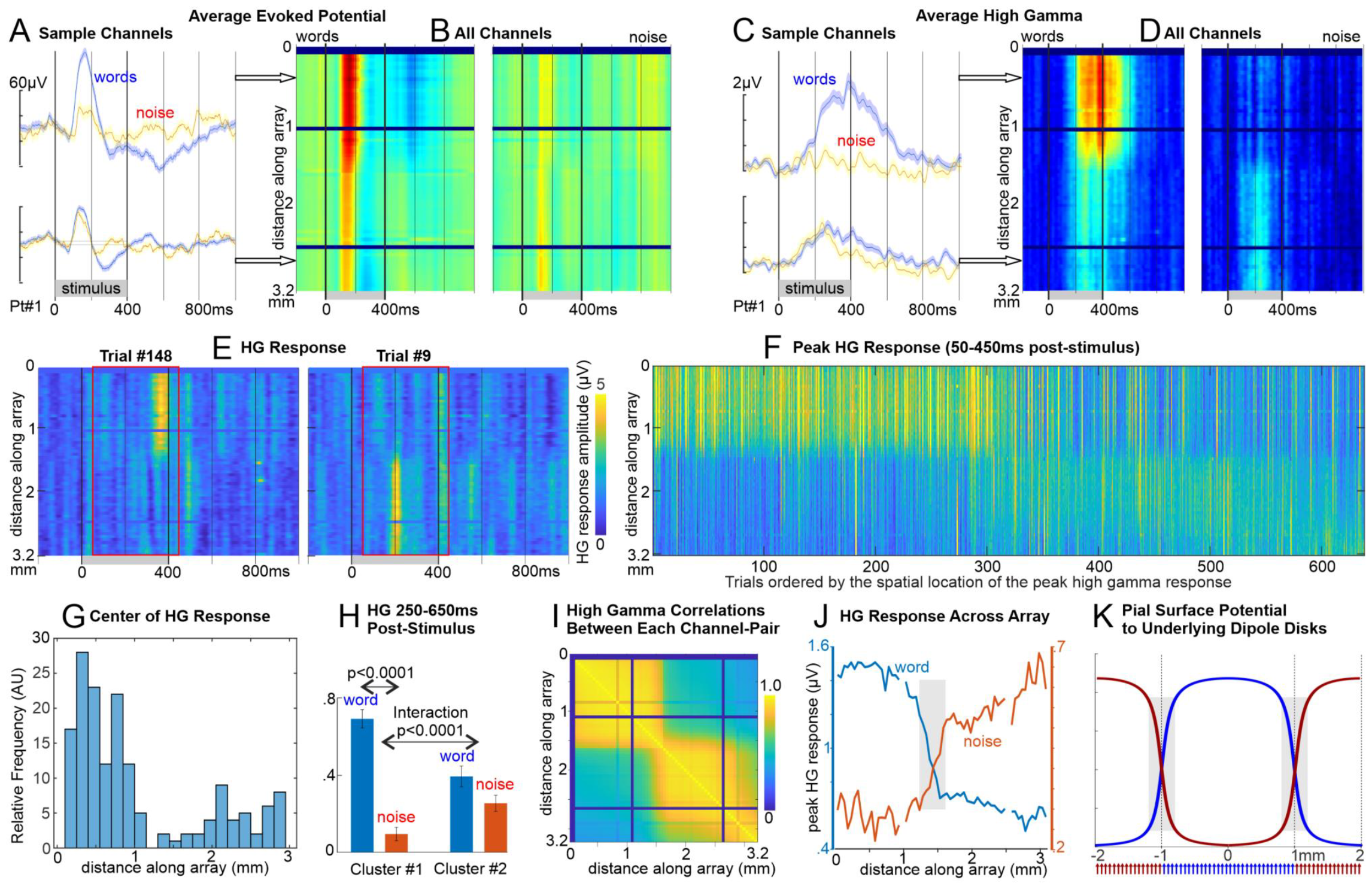
Selective responses to words. **A**. Averaged evoked potential from 2 sample channels shows greater selectivity in upper channels, with **B**. an abrupt change in response midway through the array. **C, D**. Same as A, B but analytic amplitude high gamma (70-190Hz) averaged across trials. Note that the upper channels have very consistent responses, as do the lower, suggesting two distinct functional modules. **E**. As in C, but for two individual trials showing spatially-disjoint HG burst-responses. **F**. The peak responses from each trial are plotted in accordance by the spatial middle of the response. Note the highly consistent co-activation of each module across trials. **G**. A histogram of the spatial distribution of gamma responses shows the bimodal distribution. **H**. ANOVA finds a word-selective HG response in module #1 and an interaction of word with module. **I**. HG-band activity in the first ∼25 contacts (spanning ∼1.25mm) were highly correlated over the entire task with each other, then after a transition zone of ∼.4mm (10-90% levels), the next 25 contacts were again highly correlated with each other but not with the first 25. Thus, the channels were organized into two groups, with very high within-group, and low out-group, correlation. **J**. Blue lines show the peak value of the HG average across all Word trials, at each contact; Red lines Noise trials. Note the roughly flat plateau in the modules and a sharp transition zone. **K**. The surface potential produced by alternating 2mm diameter discs of dipoles placed 100µm below the surface. The modeled curves are similar to the empirical data in panel G, suggesting possible mosaic organization of functional domains in human association cortex. The gray bars are 400µm wide and span the distance over which the curves pass from 10% to 90% of maximum, marking the transition zone. Note that the 3.2mm of the pial surface spanned by this 2×64 array at 50µ pitch would sample activity from 2 or 3 modules with ∼1.7mm diameter. Similar results from other patients are shown in Supplementary Figures 1-4.

Co-active modules were also identified in the larger arrays with Non-Negative Matrix Factorization of gamma-band (70-190Hz) analytic amplitude recorded over the entire task period (Fig. 3AE). Since the algorithm has no access to spatial information, channels are spatially co-localized because of functional similarity in signals. HG band phase-locking is very high and consistent within modules and drops rapidly at their boundaries (Fig. 3CD). The average diameter of modules with significant selectivity for words of ∼1.7mm was roughly consistent across the 3 patients, 4 recordings, and 27 modules (mean±stdev height: 1.73±0.64mm; width: 1.73±0.59mm).

**Fig 3.**
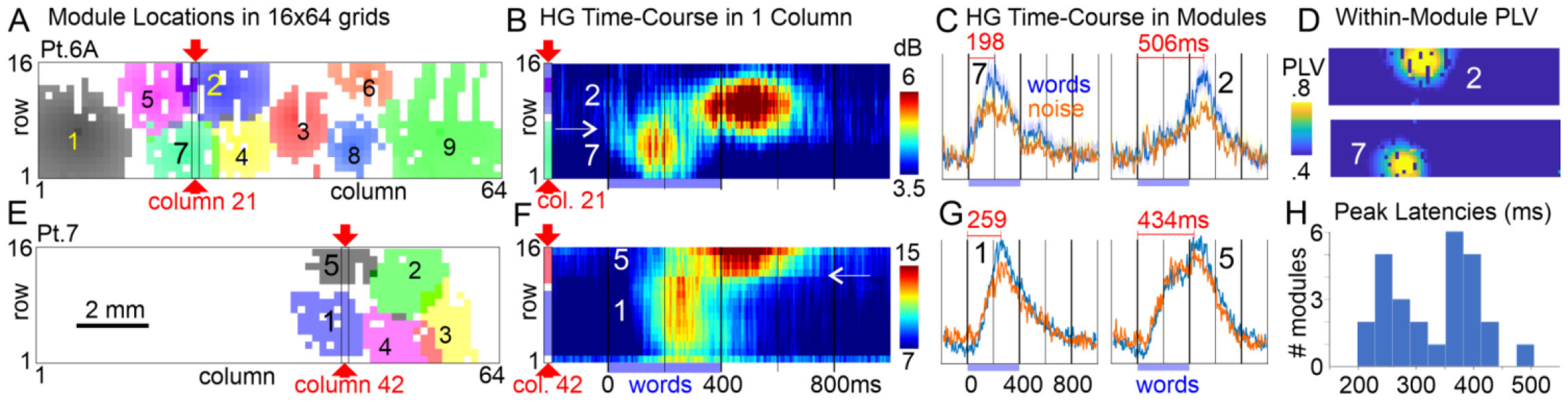
Discrete functional modules evoked by CV syllables on STG. **A**. Modules of co-active electrodes at 200µm pitch in 16×64 arrays identified with Non-Negative Matrix Factorization of HG amplitude recorded over the entire task period, in the first placement in patient 6. Module boundaries are at 2 stdev above the mean of the entire array. **B**. HG amplitude recorded from column 21 of placement 6A, passing through module 7 which peaks at ∼200ms after stimulus onset, and module 2 which peaks at ∼500ms. Columns are marked with red arrows in panel A. Note that adjacent contacts belonging to different modules have completely different time-courses of activation (→). **C**. Average HG amplitude to words and noise for modules 2 and 7. **D**. PLV between HG at the center contact of each module and the entire array. **E-G**. Same as A-C but for a traverse in column 42 of modules 1 and 5 in patient 7. Again, these modules have different latency peaks (∼250 and ∼450ms), with a sharp transition between response profiles (←). **H**. Histogram of peak module response latencies shows two distinct peaks. Similar results from other patients are shown in Supplementary Figures 5-7.

Compared to the 100ms pre-stimulus baseline, HG from 50-450ms in all 27 modules was significantly greater to the stimuli (p<.02, 2-tailed paired t-test, FDR-corrected), While 18 of 27 modules responded preferentially to words, one responded more to noise, and eight did not significantly distinguish words and noise (p<.02, 2-tailed paired t-test, FDR-corrected). Examination of contacts at module boundaries revealed a sharp transition between adjacent contacts separated by 200μm (Fig. 3BC). Peak response times in the modules were bimodal, with 13 peaking below 350ms and 14 above (Fig. 3FGH). The histogram of peak latencies was found to be marginally more parsimoniously modeled by a mixture of two gaussians than by one (Akaike information criterion = 314.08 vs 314.93) with means of 252 and 386ms. The first phoneme of each stimulus was a consonant and the second a vowel, and the latency between the average onsets of the consonants and vowels were similar to those of the early and late module response peaks. Additional recordings will be needed to test the hypothesis that consonants and vowels are encoded in adjacent sharply delineated modules.

Based on these findings we can estimate that each typical 3-4mm diameter clinical electrode would record from 4-8 modules, and at 10mm spacing would miss ∼85% of the modules. Conversely, ∼250μm diameter contacts at ∼1mm pitch would result in recordings from all modules, in most cases without significant crosstalk. Although these estimates are very rough, and assume that the modular organization exists in other association areas and are the same size, they suggest that finer sampling may reveal important clinical^16^ and basic information, and provide the approximate spatial sampling necessary in language prostheses for detecting all functional units.

In conclusion, high resolution recordings from STG provide evidence for modular organization of functional units having: (1) roughly circular shapes with ∼1.7mm diameter; (2) sharp edges with immediately adjacent modules having contrasting response time-courses and stimulus correlates; (3) roughly uniform size; and (4) continuous mapping over the sampled cortical surface forming a mosaic. Although these characteristics are consistent with a columnar organization, convincing demonstration would require oblique laminar recordings identifying a sharp transition at depth. Modular organization has also been seen for functional units with fMRI, for example for faces, but those are typically irregular ∼5-20mm patches, and do not ‘pave’ the cortex with multiple modules with sharp transitions^17^. It is possible to tile the entire cortical surface with ∼128,000 1.7mm diameter modules, each with ∼96,000 neurons. Investigation of whether such modules exist in other association cortices, their dimensions, and information-processing-activities of modules may inform multiresolution models of cortical function.

## Supplementary Information

### Online Methods

#### Acute Intraoperative Setting

When a tumor or lesion is near eloquent areas, neurosurgeons may elect to perform a resection procedure while the patient is awake, primarily using local anesthetic and systemic analgesics while minimizing sedation^15^. This technique allows the surgeon to closely monitor brain function during surgery, ensuring maximum safe resection while avoiding damage to important areas. This provides a unique opportunity for research on functional cortex while the patient is awake and interactive. Awake craniotomies, however, are relatively infrequent surgeries, and the potential time for research is limited to 20 minutes. Patients gave their informed consent to the microgrid recordings under a protocol approved by the local Institutional Review Board (OHSU IRB # 19099).

*Microarrays*^18,19^ have ultra-low impedance 30μm diameter PEDOT:PSS (typical 1kHz impedance magnitude of 38kΩ) and Pt nanorod contacts (typical 1kHz impedance magnitude of 25.5 kΩ) on a 7μm thick parylene C substrate that conforms to the pial surface, further facilitating high quality recordings. The fabrication and testing of these microarrays have previously been reported^18-20^. All devices in the experimental portion are powered via battery to avoid risk of ground loops or mains flow to the patient^21^.

*LFP pre-processing* is performed in MATLAB 2019b and LFPs inspected visually using the FieldTrip toolbox^22^. 60Hz and harmonics are notch filtered (zero-phase).

#### Time-frequency analyses

Average time-frequency plots of the ripple event-related spectral power (ERSP) are generated from the broadband LFP using EEGLAB^23^. Event-related spectral power is calculated from 1Hz to 500Hz with 1Hz resolution with ripple centers at *t*=0 by computing and averaging fast Fourier transforms with Hanning window tapering. Each 1Hz bin is normalized with respect to the mean power at -2000 to -1500ms and masked with 2-tailed bootstrapped significance with FDR correction and *α*=0.05.. Event-related phase amplitude coupling (ERPAC) is calculated using an open-source MATLAB package^24^.

*Non-negative matrix factorization (NNMF)* of high gamma (70-190Hz) analytic amplitude recorded over the entire task period was used to module the channels based on signal similarity without regard for spatial colocalization of channels^25^. The optimal number of factors for NNMF was selected by picking the approximate nadir using cross-validation; one by one, a single element is left out of the data set, and the prediction error is measured for each possible number of modules^26^. Once the optimal number of modules is defined, for each module the algorithm creates a map by assigning weights to each channel. Modules are defined by spatial colocalization of highly weighted channels; the threshold for channel inclusion is defined as weights three standard deviations above the mean for all channels for that module. Colocalized modules of channels are subsequently refined using phase-locking value, wherein the spatial center of each module was identified, and phase-locking calculated for each channel relative to the center channel. To display the boundaries of modules, a threshold value of PLV is picked that minimizes the overlap between modules.

#### Phase-locking value

PLV is an instantaneous measure of phase-locking^27^, which unlike coherence is not affected by shared amplitude modulation. PLV time courses are computed using the analytic angle of the Hilbert transformed 70-100Hz bandpassed (zero-phase shift) signals of each channel pair when there are at least 40 co-ripples with a minimum of 25ms overlap for each, and their significance calculated using FDR correction and a null distribution^28^.

#### Statistical analyses

HG relative to behavioral events are compared to an appropriate baseline control period, and between task conditions. To determine if a module was responsive to the stimuli, the Hilbert analytic amplitude from 70-190Hz was averaged on each trial in a between baseline period (100ms before stimulus onset) and in the active period (50-450ms post-stimulus onset). A 2-tailed paired t-test was performed using pairs of values from each trial. To determine if a module was differentially responsive to words versus noise, a 2-tailed t-test was performed comparing the average value in the active period between word and noise trials. All statistical tests are evaluated with α=0.02, FDR-corrected across all patients and modules^29^. Patient 7 showed responses to both words and noise but they were not significantly different, possibly due to residual anesthesia that was applied during the preceding craniotomy, which was observed behaviorally as slow responses and intermittent confusion. Patient 7’s five modules failed to distinguish between words and noise.

#### Model

The surface potential produced by alternating 2mm diameter discs of dipoles (5017 at 25μ pitch, each 10e-6 μAmm) placed 100μm below the surface in an infinite homogeneous isotropic medium (0.33 S/m). Estimates of number of modules and neurons per module based on total cortical surface area from ref^30^ and total cortical neurons in gray matter from ref^31^.

#### Task

A wide range of Consonant-Vowel syllables were presented to provide diverse activation of STG. Consonants were drawn equally from 4 categories following linguistic categories: Obstruent-Plosive (d, g), Obstruent-Fricative (s, z), Sonorant-Liquid (l, r), and Sonorant-Nasal (m, n). Vowels are drawn equally from 3 categories: Type 1 (u, aɪ, ɔ), Type 2 (ə, I, ε), and Type 3 (eɪ, æ, aʊ). Each possible CV combination is equally represented in the total stimulus set, spoken once by a male voice and once by a female. Each presented sound lasts ∼450 ms, and the SOA (stimulus onset asynchrony) was individually adjusted but usually ∼1100ms. Stimuli were presented using Presentation (Neurobehavioral Systems) in ∼6min blocks of 288 trials, each stimulus occurring once. The task was to detect CV syllables which are words.

#### Noise-Vocoding

For every CV syllable a matching noise-vocoded stimulus was constructed taking the existing biphoneme stimuli and creating a 6-band stimulus in which white-noise was multiplied by power in each of the bands to create a matched set of auditory stimuli with identical time-varying spectral acoustics^32,33^. Noise-vocoded stimuli preserve temporal envelope cues in each spectral band, providing a control for the sensory characteristics of speech, but the spectral detail within each band was degraded.

---

Figures 1 through 5 show recordings from the 2×64 channel linear array, at 50 μm pitch. Data from only one of the 2 channels at each location in the long dimension are presented for clarity; the other channel was high similar in all cases where both were technically sound. Solid, dark blue pixels indicate channels without usable signal.

**Supplementary Figure 1:**
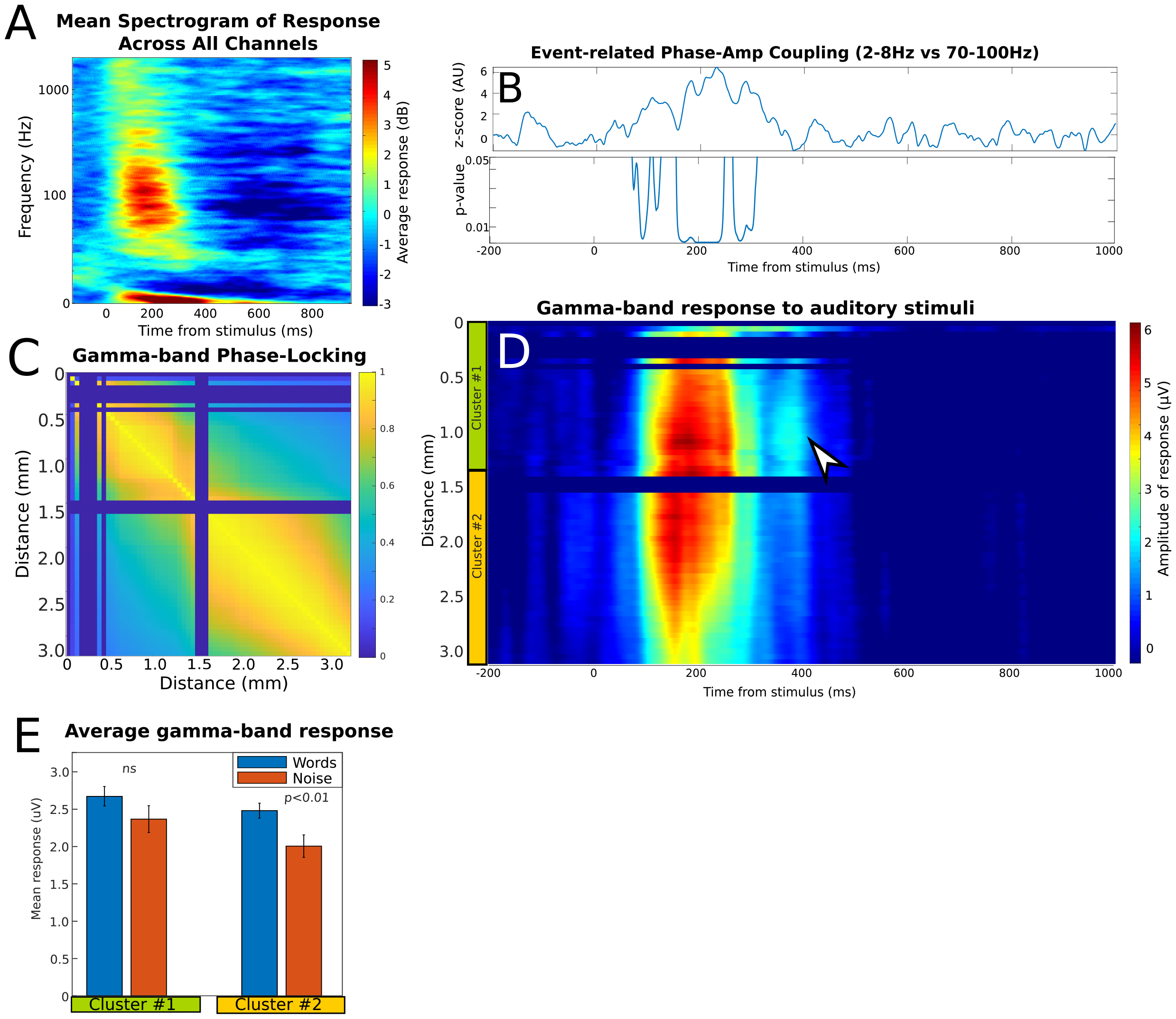
Spectral and spatiotemporal responses for the linear array in patient #4. (A) The mean event-related spectrogram of all trials shows a change in both the high gamma-band and low frequency after the onset of stimuli. (B) Event-related phase-amplitude coupling (ERPAC) (*1*) between 2-8Hz and 70-100Hz shows a stimulus-related link in response for all channels, as measured with the z-score for the ERPAC (top). FDR-corrected p-values (bottom) shows that the time points of statistical significance for the ERPAC are in the first few hundred milliseconds after the onset of the auditory stimuli. (C) A matrix of phase-locking values (PLV) between all channels shows two distinct regions of high phase-locking within but not between regions. (D) On the spatiotemporal response map, the entire electrode consistently responds to auditory stimuli, but two distinct clusters of channels are identified that factorize together (green bar, yellow bar). Close inspection shows a late temporal response in cluster #1 that separates it from cluster #2 (arrowhead). The same channels that cluster together in (D) show a high-degree of phase-locking in the gamma band (C). (E) The mean gamma-band response in the 50 – 450 ms after onset of stimuli for the clustered channels from (D) shows a stronger response to words than noise stimuli, which is only significant with cluster #2. Supplementary Figures 1 through 4 show recordings from the 2×64 channel linear array, at 50 μm pitch. Data from only one of the 2 channels at each location in the long dimension are presented for clarity; the other channel was high similar in all cases where both were technically sound.

**Supplementary Figure 2:**
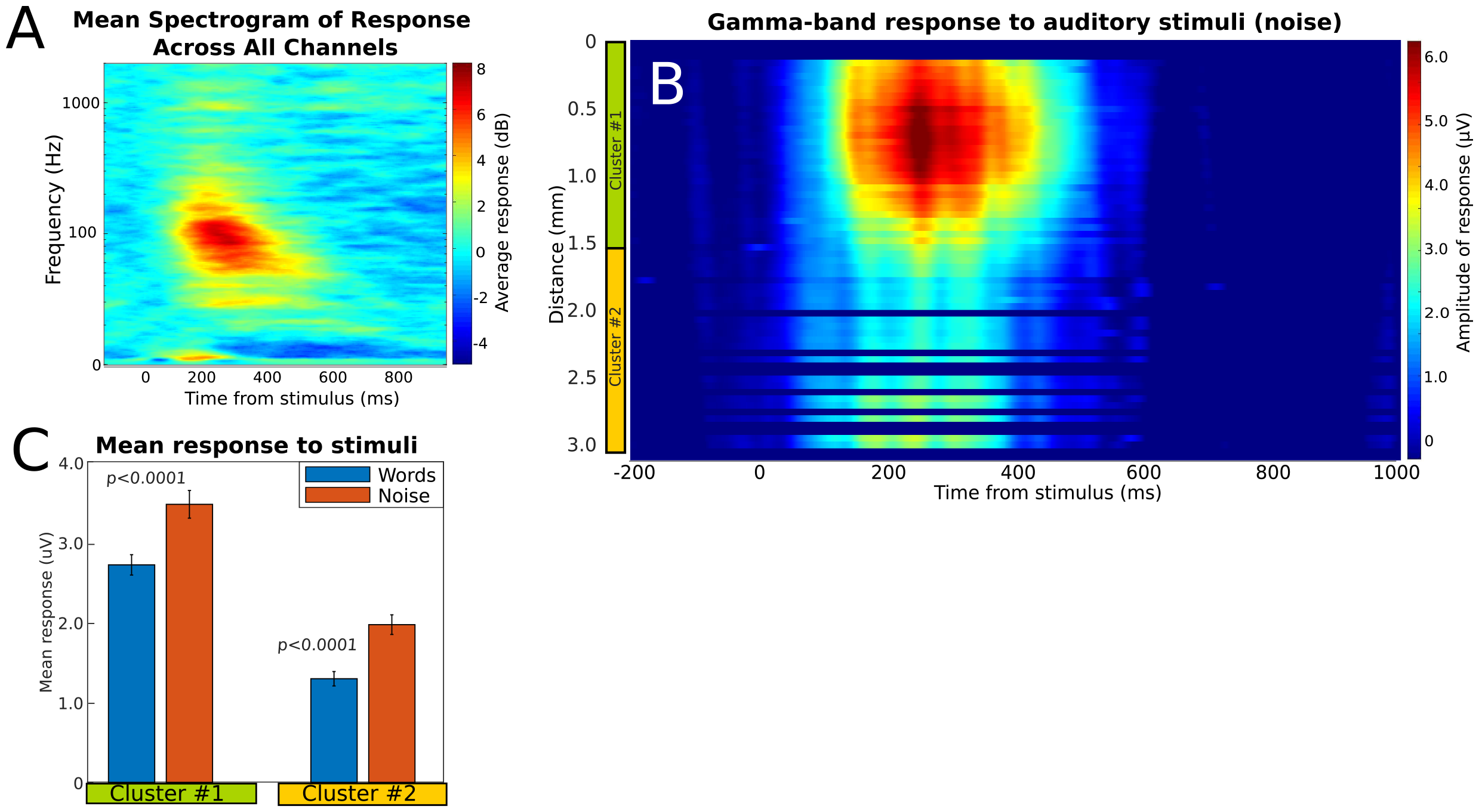
Spectral and spatiotemporal responses for linear array in patient #8, recording from non-dominant pSTG. (A) The mean event-related spectrogram of all trials shows an event-related change in both the high gamma-band and low frequency after the onset of stimuli. (B) On the spatiotemporal response map of gamma activity, increased activity was observed in the few hundred milliseconds after onset of auditory stimuli at time zero. Two distinct spatial segments are identified using factorization to cluster the channels (green bar, yellow bar). (C) Quantifying the mean gamma-band response in the 50 – 450 ms after onset of stimuli for the clustered channels shows a stronger response to noise than word stimuli for both clusters of channels.

**Supplementary Figure 3:**
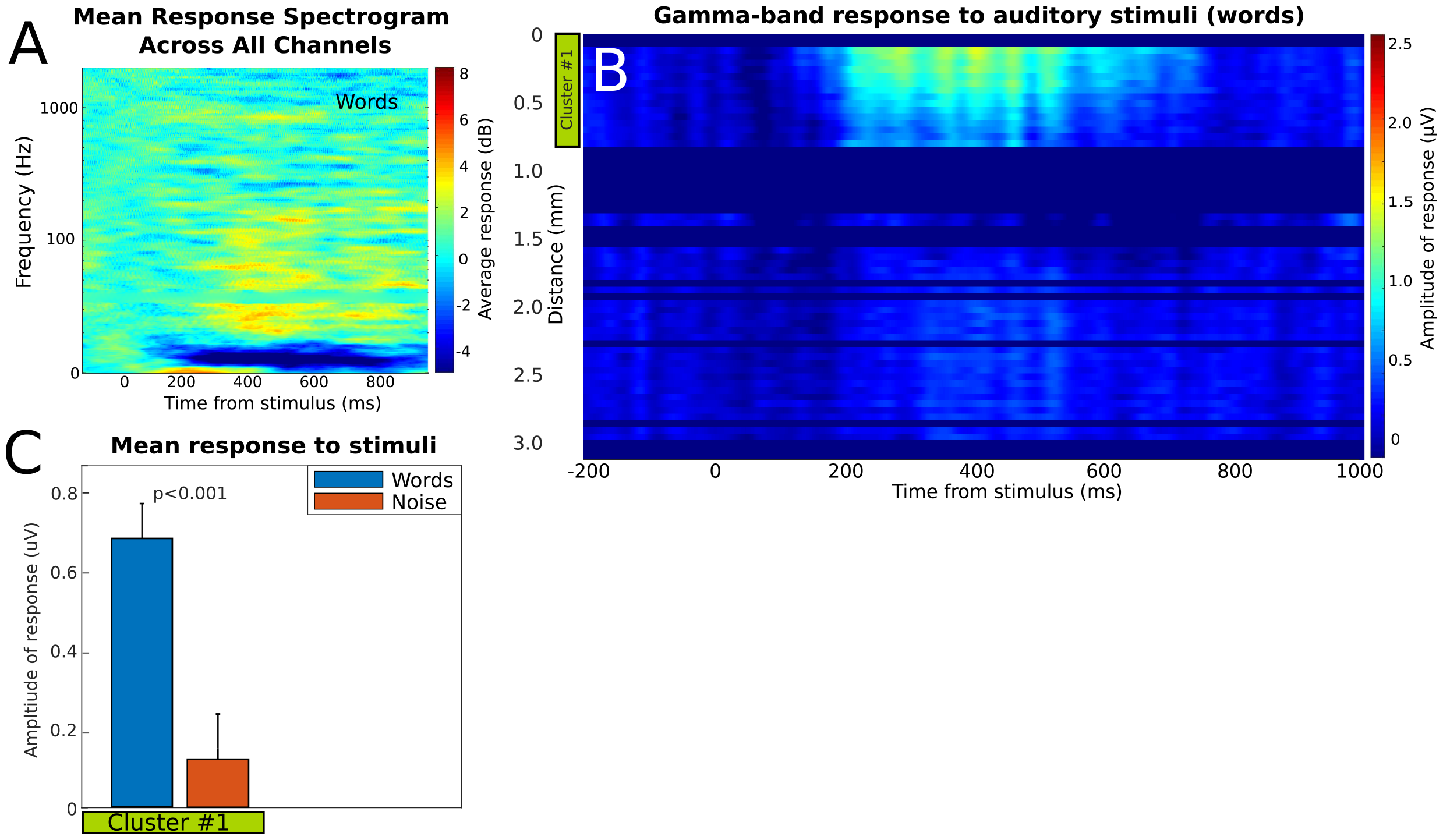
Spectral and spatiotemporal responses for linear array in patient #3. (A) The mean event-related spectrogram of all trials shows an lower amplitude of increase in both the high gamma-band and low frequency than observed in other patients. (B) Similarly, on the spatiotemporal response map of gamma activity, increased activity was only observed in one segment for in the few hundred milliseconds after onset of auditory stimuli. Only one spatial segment was identified using factorization (green bar). (C) Quantifying the mean gamma-band response in the 50 – 450 ms after onset of stimuli for the clustered channels shows a stronger response to words than noise stimuli.

**Supplementary Figure 4:**
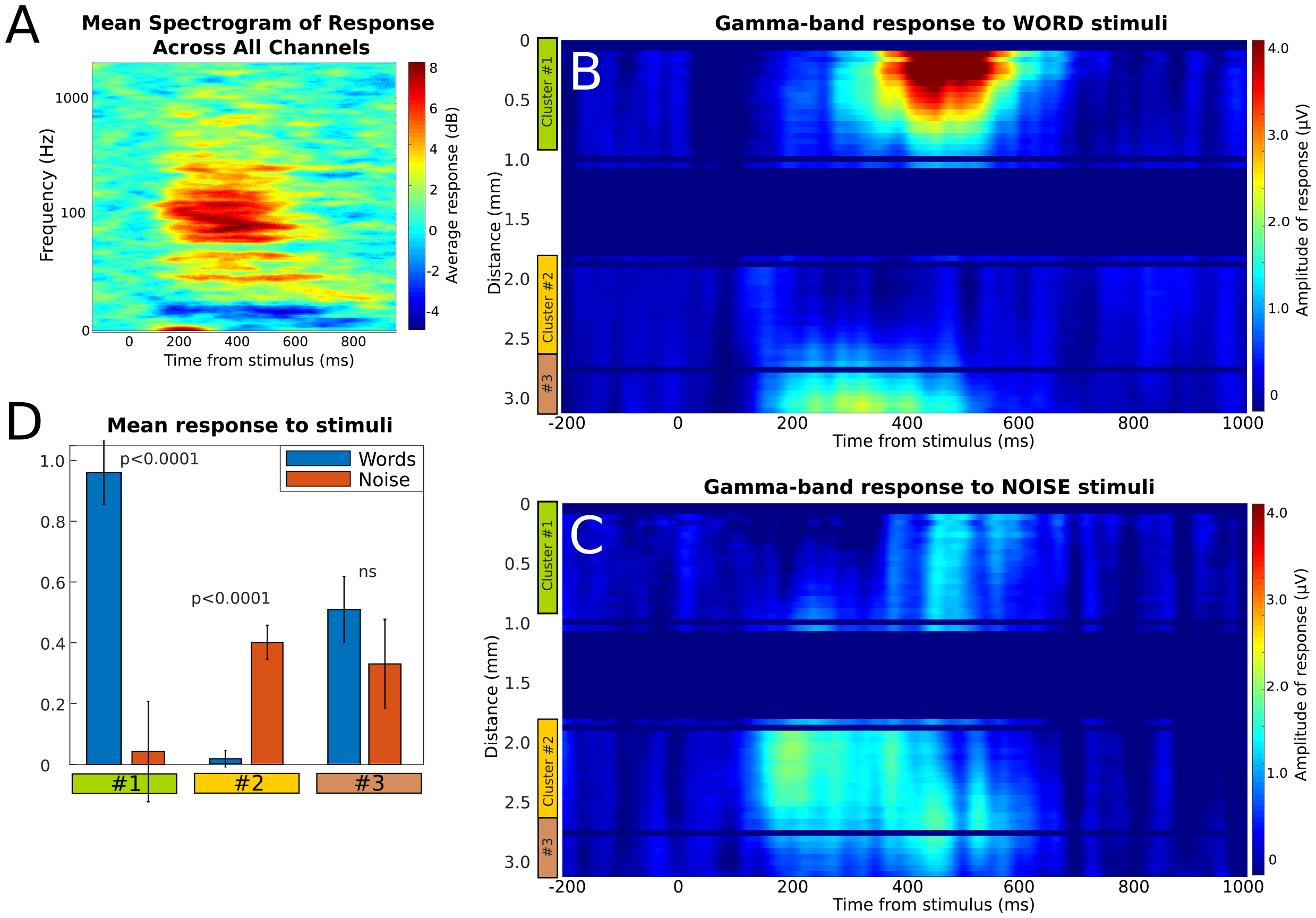
Spectral and spatiotemporal responses for linear array in patient #2. (A) The mean event-related spectrogram of all trials shows a change in both the high gamma-band and in low frequencies after the onset of stimuli. (B) On the spatiotemporal response map of gamma activity for words, spatially segregated increase activity was observed in the few hundred milliseconds after onset of auditory stimuli at time zero. Three distinct spatial segments are identified using factorization to cluster the channels, two of which appear to preferentially respond to word stimuli (green bar, brown bar) while one segment appears to preferentially respond to noise stimuli (yellow bar). (C) Quantifying the mean gamma-band response in the 50 – 450 ms after onset of stimuli for the clustered channels shows a stronger response to words than noise stimuli for one of the clusters of channels (green cluster), a stronger response to noise for one cluster (yellow cluster), and an indeterminate response for one cluster (brown cluster).

**Supplementary Figure 5:**
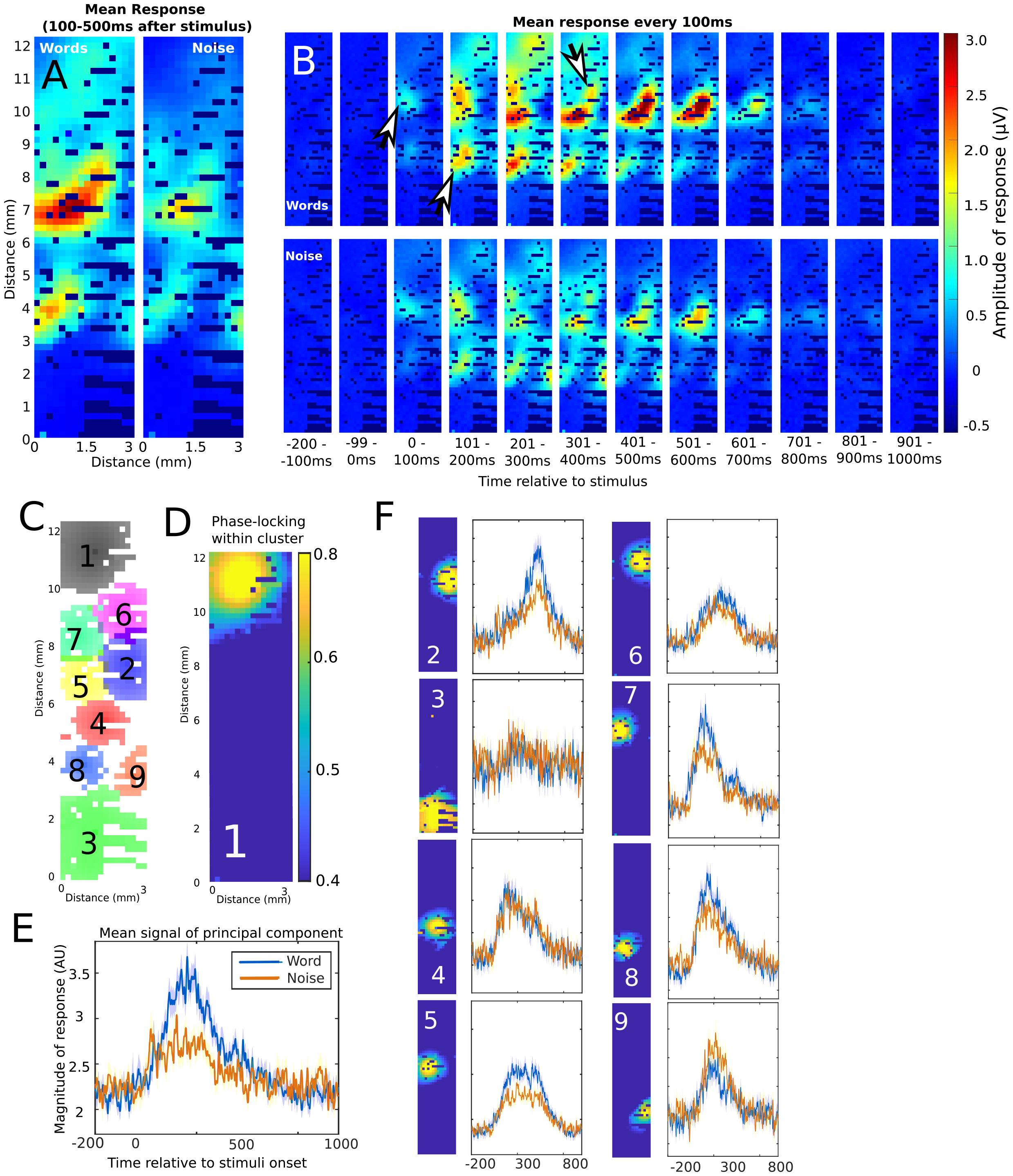
Spatiotemporal maps of gamma-band responses for Patient #6A, using the 1024-channel electrode array (16×64 with 200 μm spacing). (A) The mean gamma-band response between 100 ms and 500 ms after the onset shows multiple poorly demarcated regions increased signal. (B) Finer temporal discrimination is used to view gamma-band responses every 100 ms, which shows distinct spatial regions of activation. The spatial regions have distinct temporal onset and offset. Arrowheads indicate spatial regions with unique temporal activation times. (C) A spatial map of the 16×64 channel electrode with the results of factorization overlaid. Factorization identified nine distinct, spatially adjacent regions. (D) For an individual cluster of channels (Module #1), the phase-locking value for each cluster shows a high degree of concordance for spatially adjacent channels with rapid drop off. (E) The mean signal of the principal components of Module #1 shows the temporal and feature preference for the underlying channels. The mean response is calculated using the average response relative to the start of the stimuli for all trials. (F) The other modules identified by factorization similarly show distinct temporal properties in terms of average onset, offset, and response features.

**Supplementary Figure 6:**
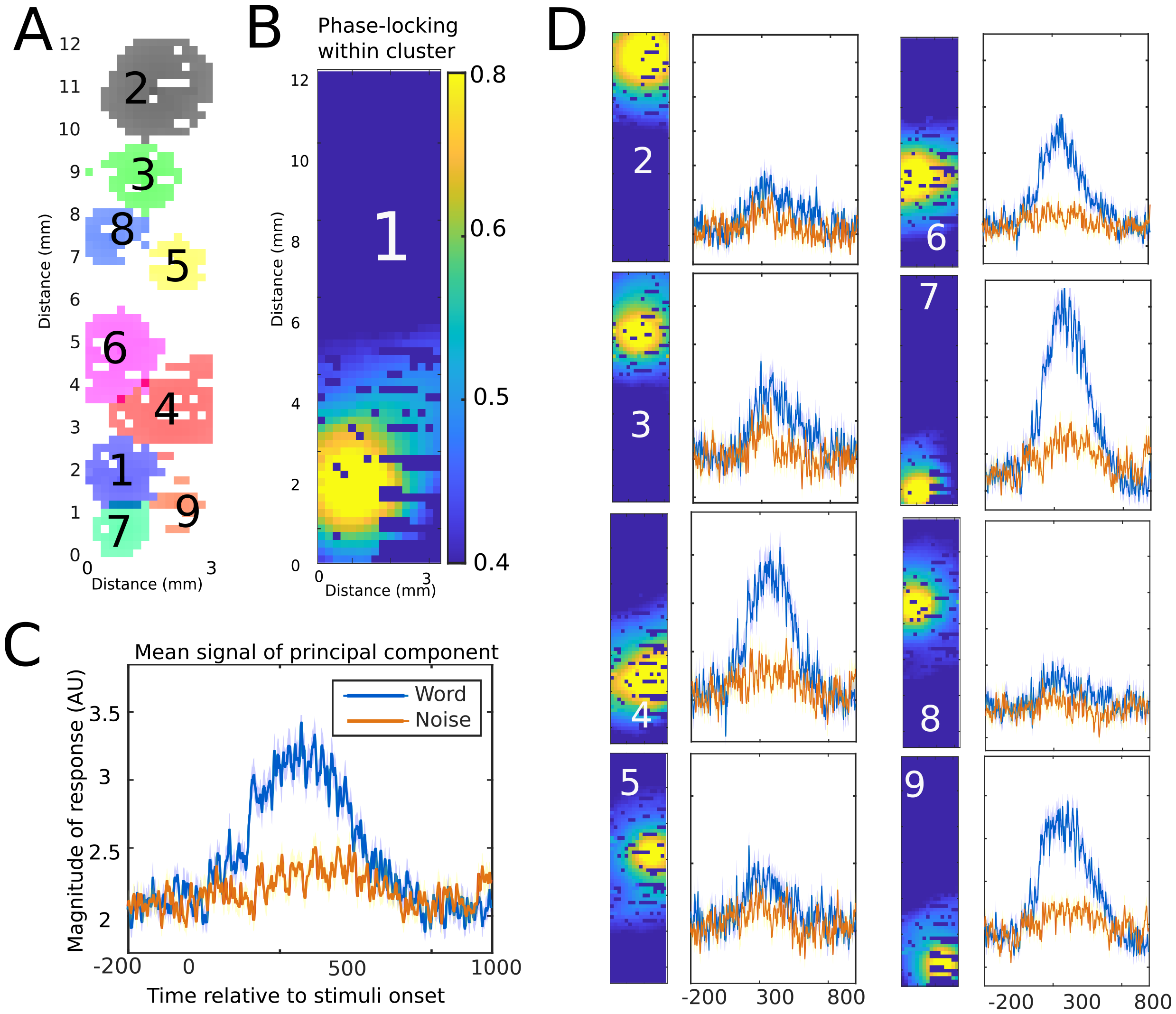
Spatiotemporal maps of gamma-band responses for Patient #6B, using the 1024-channel electrode array (16×64 with 200 μm spacing). A) A spatial map of the 16×64 channel electrode with the results of factorization overlaid. Factorization identified nine distinct, spatially adjacent regions. (B) For an individual cluster of channels (Module #1), the phase-locking value relative for cluster shows a high degree of concordance for spatially adjacent channels with rapid drop off. (C) The mean signal of the principal components of Module #1 shows the temporal and feature preference for the underlying channels. The mean response is calculated using the average response relative to the start of the stimuli for all trials. (D) The other modules identified by factorization similarly show distinct temporal properties in terms of average onset, offset, and response features.

**Supplementary Figure 7:**
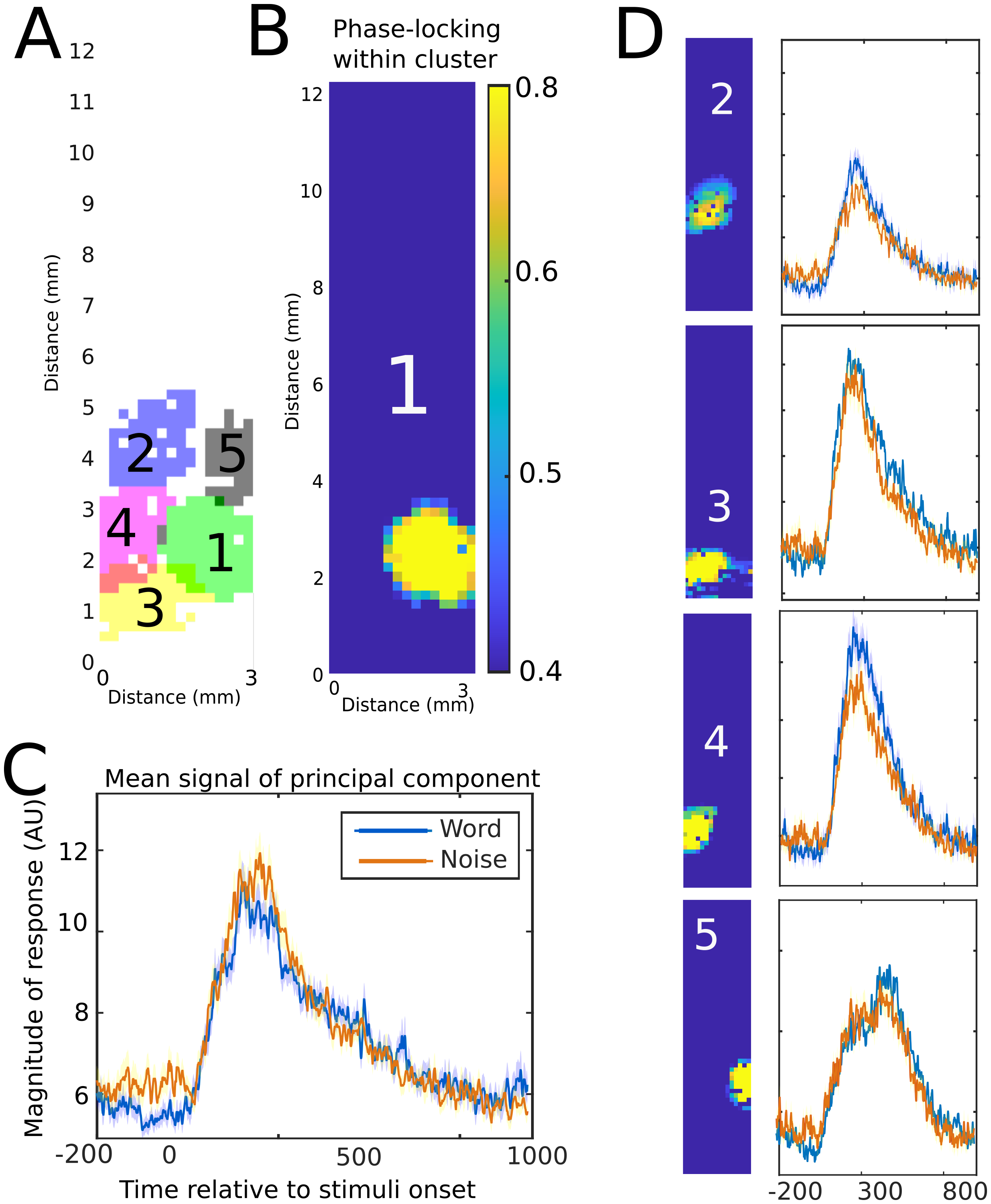
Spatiotemporal maps of gamma-band responses for Patient #7, using the 1024-channel electrode array (16×64 with 200 μm spacing). A) A spatial map of the 16×64 channel electrode with the results of factorization overlaid. In this patient, factorization identified five distinct, spatially adjacent regions. (B) For an individual cluster of channels (Module #1), the phase-locking value relative for cluster shows a high degree of concordance for spatially adjacent channels with rapid drop off. (C) The mean signal of the principal components of Module #1 shows the temporal and feature preference for the underlying channels. The mean response is calculated using the average response relative to the start of the stimuli for all trials. (D) The other modules identified by factorization similarly show distinct temporal properties in terms of average onset, offset, and response features.

